# Multiple retrospective analysis of survival and evaluation of cardiac death predictors in a population of dogs affected by degenerative mitral valve disease in ACVIM class C treated with different therapeutic protocols

**DOI:** 10.1101/2020.01.13.904102

**Authors:** Mara Bagardi, Chiara Locatelli, Annamaria Zanaboni, Alberto Galizzi, Dario Malchiodi, Paola G. Brambilla

**Author notes:** These authors contributed equally to this work (Mara Bagardi, Chiara Locatelli). Corresponding author (MB).

## Abstract

Clinical records of dogs with spontaneous degenerative mitral valve disease (DMVD) with clinical signs related to congestive heart failure (CHF) that had been recruited during routine clinical practice, between 2001 and 2018 at the Cardiology Unit of the Veterinary Teaching Hospital (University of Milan) were included in this retrospective cohort study. Baseline echocardiographic data were evaluated. Median survival times (MSTs) were calculated. Data on therapeutic treatment, ISACHC or ACVIM classes were reviewed based on the inclusion period and type of endpoint (i.e. cardiac death or death for other causes). The main goal of this data review was to retrospectively evaluate 259 clinical records of subjects belonging to ACVIM C class examined between 2001 to 2018 together with the 202 examined between 2010 to 2018. The MSTs of these subjects was 531 d (2001-2018) and 335.5 d (2010-2018), respectively. Univariate survival regression analysis for subjects included from 2010 to 2018 showed the following variables as being significantly related to cardiac death (CD): LA/Ao ratio (HR 2.754, p=0.000), E wave (HR 2.961, p=0.000), E/A ratio (HR 1.372, p=0.000), EDVI (HR 1.007, p=0.000), ESVI (HR 1.012, p=0.026), Allo(d) (HR 4.018, p=0.000) and Allo(s) (HR 2.674, p=0.049), age (HR 1.006, p=0.009) and PH severity (HR=1.309, p=0.012). Multivariate analysis, adjusted for age, showed that the only variable that determined a statistically significant difference in MST was PH severity (HR 1.334, p=0.033). The type of therapeutic treatment within this class was not significant for the MST of the subjects.

## Introduction

Degenerative mitral valve disease is the most common heart disease in middle-old aged and medium-sized dogs. It is characterized by a slow progression over years, and in many affected dogs, because of the age of onset, does not always progress to clinical signs of congestive heart failure (CHF) [1,2,3]. Some authors have reported a survival time of between 5 to 14 months after the onset of clinical signs [4,5]. Although DMVD has been studied for more than 40 years, its treatment remains a challenge for the clinician.

Many advances have been made regarding diagnosis, imaging, medical and surgical therapy, however few studies have evaluated the natural history and prognostic factors of the disease in dogs [3,6,7]. Furthermore, to the best of our knowledge, no studies have evaluated MST in relation to different types of multiple therapeutic treatment within the same ACVIM C class severity class.

The aim of DMVD treatment is to modulate hemodynamic and neurohormonal disorders, including high venous pressures, reduce the systolic function, activate the sympathetic and renin-angiotensin-aldosterone systems, and also release cytokines and vasopressin [1,2]. Treatment of CHF due to DMVD consists of a diuretic (loop diuretic, ++ furosemide) and additional agents (angiotensin-converting enzyme inhibitors, inodilators and aldosterone receptor antagonists) [8,9]. Angiotensin-converting enzyme inhibitors (ACE-Is), benazepril or enalapril, combined with furosemide improve the quality of life [4,10]. The administration of pimobendan, an inodilator, has been evaluated in dogs with CHF due to DMVD [5,11].

Several studies have compared symptomatic dogs with DMVD that have received pimobendan and furosemide versus dogs treated with an ACE-I (ramipril or benazepril) and furosemide [5,12]. Dogs that received pimobendan and furosemide showed an improvement in clinical signs and quality of life, and a reduction in the probability of developing an adverse cardiac event [5,12]. VetSCOPE and QUEST reported that dogs receiving pimobendan survived longer than dogs that did not [5,13]. Improved survival and reduction of risk for a cardiac event have also been shown in dogs affected by DMVD and CHF treated with spironolactone [14,15].

The main goal of this study was to retrospectively investigate the survival time of a population of dogs affected by DMVD belonging to ACVIM class C and treated with different combinations of drugs including furosemide, ACE-I (benazepril or enalapril), pimobendan, and spironolactone.

The effects of the different therapeutic protocols on MST, and the prognostic value of the echocardiographic variables were also evaluated.

## Materials and methods

### Study design

This is a retrospective cohort study. The clinical records of dogs affected by DMVD examined at the Cardiology Unit of the Veterinary Teaching Hospital (University of Milan) between 2001 and 2018 were reviewed. Owner consent was routinely requested before the first examination of each dog. It was not necessary to obtain authorization from the Ethics Committee because this is a retrospective study carried out on data collected in subjects routinely brought to a clinical examination by the owners.

From the beginning of the study to 2009, the admitted dogs were classified according to ISACHC classes [16], and from 2010 to 2018 according to the ACVIM classification [9]. In order to compare the subjects, a univocal classification was needed, and the patients classified in ISACHC classes II, IIIa and IIIb were reallocated to ACVIM class C. Data obtained from clinical records from 2001 to 2018 were then statistically analysed.

Later, in order to avoid inclusion bias due to the reallocations, statistical analyses were performed on a selection of subjects belonging to ACVIM class C, recruited from 2010 to 2018.

Specific attention was focused on the therapy changes, as well as detailed information on any change in ACVIM class and the causes of death reported in clinical records of subjects included from 2010 to 2018 were studied.

### Inclusion criteria

The clinical records were selected according to the following inclusion criteria: complete clinical findings including signalment, history, physical examination, thoracic radiographs, electrocardiogram (ECG) and a diagnosis of DMVD ACVIM class C based on echocardiographic, and Doppler evaluation associated with clinical signs (increased resting respiratory rate, cough, dyspnoea, ascites) [17]. In all subjects the presence and severity of PH were evaluated, based on the TRV. The PH was classified as reported in the literature [18,19]. The accepted administered drugs were a combination of diuretics (furosemide), ACE-I (benazepril, enalapril and ramipril), inodilator (pimobendan), and spironolattone. The therapeutic protocol applied was clearly reported on the clinical record from the first to the last examiner, as well as whether the owner was willing to be interviewed by telephone. In this study all genders, weights and breeds were included, except for Cavalier King Charles Spaniel [8,20,21]. The clinical records of subjects for whom cardioactive therapy had previously been set up were also included.

### Exclusion criteria

Clinical records of subjects affected by any other heart disease apart from DMVD and/or with concurrent congenital heart disease and acquired cardiovascular disorders that could affect the mitral valve or its functions (bacterial endocarditis, myocardial disease, arrhythmias) were excluded. Subjects with primary hypertension were not included in the study [22]. Incomplete clinical records or with missing information on the therapeutic protocol adopted were also excluded.

### Echocardiography

Echocardiographic examinations were performed on conscious dogs by specialists in cardiology, and in accordance with the guidelines of the American Society of Echocardiography using the leading edge-to-leading edge method for M-mode measurements and Hansson’s method for 2-dimensional (2D) measurements of the left atrial (LA) and aortic root (Ao) diameters [23,24].

### Follow-up and endpoints

A single investigator (M.B.) conducted telephone interviews with dog owners to determine the clinical outcome of each dog. For this the following information was obtained: was the dog dead or alive, had the dog been euthanized or did it die spontaneously, and reasons for euthanasia or cause of death. The date and cause of death (either spontaneous death or euthanasia) were recorded. Dogs euthanized for severe refractory heart failure were considered as cardiac-related deaths. Sudden deaths were counted as cardiac-related if no other cause of death was obvious.

Dogs still alive, dead or euthanized for reasons unrelated to cardiac disease were removed from the statistical analysis; subjects lost to follow-up were included in the survival analysis up to the last time point at which they were known to be alive and were then removed from the analysis.

The survival analysis was performed considering different end points such as death due to other causes (OC), death related to the studied heart pathology (CD – cardiac death), first and following therapy changes and moving to more advanced gravity class. The survival time in the CD group was also analysed in relation to the therapeutic scheme.

The selected clinical records had to report the date and the medications prescribed as well as any variation in therapy during the follow-up.

Median survival time (days) was calculated from the admission date to death or to the last contact with the owner (lost to follow-up). The MST of each patient was subsequently related to the combination of medicines (groups 1, 2 and 3) and the echocardiographic data (LA/Ao ratio, E wave, E/A ratio, FE%, FS%, EDVI, ESVI, Allo(d), Allo(s), TRV and PH) at the time of inclusion.

### Statistical analysis

The data obtained for the analysis were compiled on an Excel spreadsheet and then processed with SPSS^TM^ 25.0 (IBM, SPSS, USA). The statistical analysis was performed in two different steps. In the first part of the study, which included the analysis of the reallocated ACVIM C class population between 2001 and 2018, the statistical analysis was essential to verify the presence of correlations and their significance between the MST and therapy group and echocardiographic parameters such as LA/Ao ratio, E wave, E/A ratio, FE%, FS%, EDVI, ESVI, Allo(d) and Allo(s).

The analysis thus included clinical records of subjects belonging to the ACVIM C class included from 2010 to 2018 with the same inclusion criteria as the previous analysis. The prognostic CD values of echocardiographic measurements, presence and severity of PH, and the modulation of therapeutic protocols for each patient over the follow up time were estimated.

A descriptive analysis of the sample was performed in terms of mean and standard deviation or median and interquartile range (IQR) for normally or non-normally distributed variables, respectively. Differences between variables were assessed by the appropriate test (Student t-test for independent or paired normal variables, Mann Whitney U-test for independent non-normal variables, and Wilcoxon signed ranks test; and the sign test for paired non-normal variables). The therapeutic groups were compared by ANOVA and Tukey’s HSD test. Median survival times were compared by the log rank test.

The correlation between variables was investigated by the Pearson correlation or the Kendall tau test, as appropriate. The Kolmogorov-Smirnov test for independent samples was used to compare distributions.

The influence of individual physical and echocardiographic parameters on the MST was assessed by univariate and multivariate survival analysis. Correlations between variables were calculated to: (a) determine the possible predictors in the multivariate regression model, (b) select only uncorrelated variables as predictors. The hazard ratio (HR) for each variable was also evaluated.

A confidence interval (CI) of 95% was considered. The differences were considered as statistically significant with p<0.05.

## Results

### Results ACVIM C class (2001-2018)

Six hundred and thirty-one clinical records from 2001 to 2018 reported a diagnosis of DMVD with varying levels of gravity. After the reclassification (conversion from ISACHC to ACVIM), the following 259 clinical records fulfilled the inclusion criteria and were thus included: 135 (52%) intact male dogs, 29 (11%) neutered males, 60 (23%) sterilized females, and 35 (13%) intact females. The median weight was 11.07 Kg (CI=5.8-14), the median age was 11.89 (CI=10.37-13.87) years. Any breed of dog was included of which: 125 (48.2%) were mixed breed, 20.5 (7.9%) Poodle, 20 (7.7%) Yorkshire Terrier, 13 (5.1%) Dachshund, 7 (2.8%) Shi-Tzu, 6 (2.6%) Pinscher, as well as lower percentages of other breeds.

The therapeutic groups considered in the first analysis are reported in Table 1. Table 2 reports the average, median, standard deviation, minimum and maximum age, weight, LA/Ao ratio, E wave, E/A ratio, FE%, FS%, EDVI, ESVI, Allo(d) and Allo(s) of all subjects included in the analysis belonging to C ACVIM class.

**Table 1:**
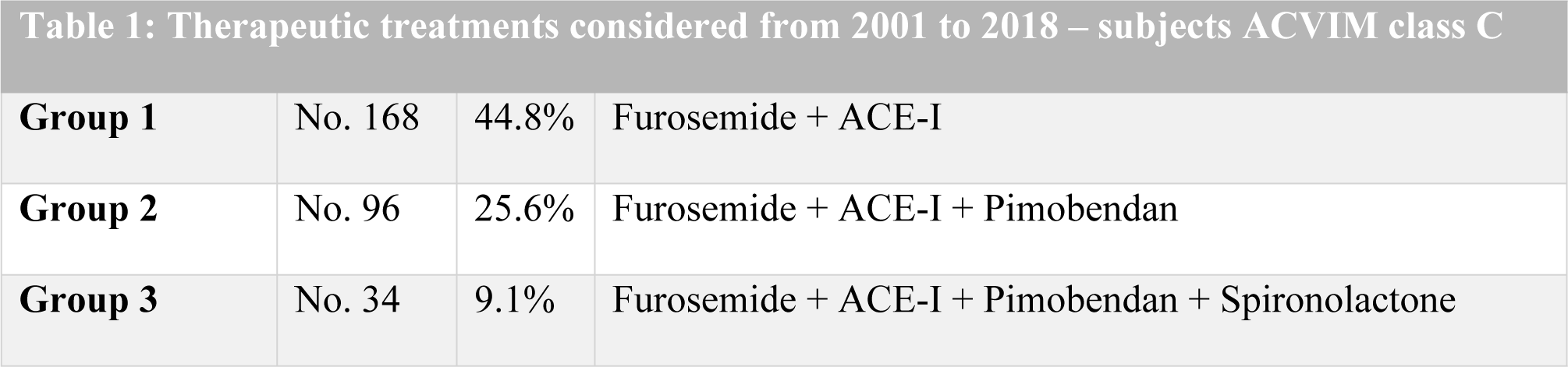
Classification of therapeutic treatments considered in the first analysis (subject in ACVIM class C from 2001 to 2018). Clinical records of dogs for whom cardioactive therapy had previously been arranged were also included.

|  |  |  |  |
| --- | --- | --- | --- |
| **Group 1** | No. 168 | 44.8% | Furosemide + ACE-I |
| **Group 2** | No. 96 | 25.6% | Furosemide + ACE-I + Pimobendan |
| **Group 3** | No. 34 | 9.1% | Furosemide + ACE-I + Pimobendan + Spironolactone |

**Table 2:** Summary of the physical and echocardiographic parameters assessed in the statistical analysis of all the dogs included in the first analysis belonging to ACVIM class C. All these parameters had a normal distribution. SD: standard deviation.

| Table 2: Physical and echocardiographic parameters of all dogs belonging to ACVIM class C (2001-2018) |  |  |  |  |  |  |  |  |  |  |  |  |
| --- | --- | --- | --- | --- | --- | --- | --- | --- | --- | --- | --- | --- |
| CLASS |  | Weight<br>(Kg) | Age | LA/Ao | E | E/A | FE % | FS% | EDVI | ESVI | Allo<br>diastolic | Allo<br>systolic |
| C | Average | 11.27 | 12.02 | 2.29 | 1.33 | 1.65 | 73.89 | 43.56 | 156.77 | 40.86 | 2.02 | 1.16 |
|  | No. | 259.00 | 179.00 | 257.00 | 127.00 | 121.00 | 255.00 | 255.00 | 257.00 | 257.00 | 258.00 | 258.00 |
|  | SD | 8.51 | 2.90 | 0.49 | 0.38 | 0.71 | 10.41 | 9.51 | 60.92 | 25.22 | 0.33 | 0.26 |
|  | Median | 8.00 | 12.49 | 2.22 | 1.36 | 1.47 | 76.00 | 44.00 | 148.02 | 36.10 | 2.02 | 1.15 |
|  | Interval | 51.00 | 17.37 | 3.10 | 2.32 | 4.04 | 65.00 | 87.00 | 377.20 | 205.45 | 2.34 | 1.86 |
|  | Minimum | 2.00 | 0.14 | 1.30 | 0.48 | 0.15 | 28.00 | 13.00 | 6.22 | 3.83 | 0.58 | 0.49 |
|  | Maximum | 53.00 | 17.51 | 4.40 | 2.80 | 4.19 | 93.00 | 100.00 | 383.42 | 209.28 | 2.92 | 2.34 |

Class ACVIM C included 259 subjects, 136 of which (52.5%) died of CD with an MST of 531 days, and 123 dogs (47.5%) were still alive at the end of the study or died of OC. The MST was 318 days (Table 3).

**Table 3:** Univariate and multivariate analysis of clinical records from 2001 to 2018. Parameters not reaching statistical significance are not reported in the table.

| Table 3: Clinical records analysis from 2001 to 2018 |  |  |  |  |
| --- | --- | --- | --- | --- |
| *Results ACVIM C class (2001-2018)* |  |  |  |  |
| ACVIM Class | No. of subjects | No. and % of<br>CD | No. and % of alive or dead<br>due to OC | MST days (CI 95%) for CD |
| C | 259 | 136 (52.5%) | 123 (47.5%) | 531 (440.812-621.188) |
| *Univariate and multivariate analysis in ACVIM class C* |  |  |  |  |
| Predictors | Univariate analysis HR (95%<br>CI) | p value | Multivariate analysis HR (95% CI) | p value |
| LA/Ao ratio | 2.473 (1.784-3.429) | 0.000 | 2.473 (1.784-3.429) | 0.000 |
| E wave | 2.560 (1.489-4.010) | 0.001 | - | - |
| E/A ratio | 1.687 (1.222-2.329) | 0.001 | - | - |
| EDVI | 1.004 (1.001-1.007) | 0.002 | - | - |
| ESVI | 1.006 (1.001-1.011) | 0.021 | - | - |
| Allo(d) | 2.197 (1.282-3.763) | 0.004 | - | - |
| Allo(s) | 2.058 (1.126-3.764) | 0.019 | - | - |
| Age | 1.080 (1.005-1.162) | 0.037 | - | - |
| Administration of spironolactone | 0.623 (0.403-0.963) | 0.033 | - | - |
| ***Analysis of the groups of treatment in ACVIM class C*** |  |  |  |  |
| Therapeutic groups | n. of subjects | n. and % of CD | n. and % of alive or death due to OC | MST days (CI 95%) |
| 1 | n. 145 | 66 (45.4%) | 79 (54.6%) | 665 (532.724-794.276) |
| 2 | n. 80 | 45 (56.2%) | 35 (43.8%) | 487 (306.392-667.608) |
| 3 | n. 34 | 25 (73.3%) | 9 (26.7%) | 447 (190.470-703.530) |
| ***Univariate analysis of the groups of treatment in ACVIM class C*** |  |  |  |  |
| Predictors | Univariate analysis HR (95% CI) | p value | Multivariate analysis HR (95% CI) | p value |
| LA/Ao ratio | 2.773 (1.978-3.880) | 0.000 | - | - |
| administration of therapy | 1.295 (1.020-1.644) | 0.033 | - | - |
| (yes/no) | 2.686 (1.571-4.594) | 0.000 | - | - |
| E wave | 1.669 (1.200-2.321) | 0.002 | - | - |
| E/A ratio | 1.005 (1.002-1.008) | 0.000 | - | - |
| EDVI | 1.006 (1.001-1.012) | 0.014 | - | - |
| ESVI | 2.821 (1.607-4.952) | 0.000 | - | - |
| Allo(d) | 2.222 (1.218-4.054) | 0.009 | - | - |
| Allo(s) |  |  |  |  |

Univariate regression analysis showed the following variables to be statistically significant: LA/Ao ratio, E wave, E/A ratio, EDVI, ESVI, Allo(d), Allo(s), Age and administration of spironolactone. All the variables analysed were statistically significant for survival and their increment was found to be related to an increase in the risk of death, except for the administration of spironolactone (HR < 1).

Multivariate analysis highlighted that the LA/Ao ratio correlated negatively to MST and significantly increased the risk of death (by 2.5 times) (Table 3).

### Analysis of the groups of treatment in ACVIM class C (2001-2018)

The therapy groups in the ACVIM class C were 1, 2 and 3 (Table 1).

Group 1 included 145 subjects, 66 (45.4%) died of CD, and 79 (54.6%) died of OC, and the others were still alive at the end of the study. Group 2 included 80 dogs, 45 (56.2%) died of CD and 35 (43.8%) died of OC or were still alive at the end of the study. Group 3 was comprised of 34 dogs, 25 of which (73.3%) died of CD and 9 died (26.7%) of OC or were still alive at the end of the study.

The MST of subjects that died of CD was 665 days in group 1, 487 days in group 2, and 447 days in group 3.

The univariate analysis revealed a positive correlation among the LA/Ao ratio, administration of therapy (yes or no), E wave, E/A ratio, EDVI, ESVI, Allo(d) and Allo(s)), and CD. The univariate analysis of the aforementioned variables was statistically significant for MST, whose increase, increased the risk of death (Table 3).

The multivariate analysis for subjects in class ACVIM C showed that only LA/Ao led to a statistically significant difference in MST, and significantly increased (2.5 times) the risk of CD, as described in Table 3.

MSTs were also evaluated together with the influence of individual parameters on the MST using univariate analysis and ANOVA tests between subjects with different types of therapy. Using Tukey HSD tests, a multiple comparison was performed between the therapeutic classes listed above. The log-rank method also showed that there was no statistically significant difference (p=0.091) between the MSTs for the CD of dogs undergoing different cardioactive therapies. Univariate analysis for subjects in the ACVIM class C with different cardioactive therapies showed that the LA/Ao ratio (p=0.000), E wave (p=0.000) and EDVI (p=0.01) led to a statistically significant difference in MST, some of which significantly increased the risk of CD, as did some of the other variables considered (Table 3).

The multiple comparisons among therapeutic classes 1, 2 and 3 executed by ANOVA were confirmed by Tukey’s HSD test. The LA/Ao ratio, E wave and EDVI were statistically significant (p<0.001) between therapeutic groups 1 and 2, 1 and 3, and not between groups 2 and 3. The statistical analysis was then carried out with the log-rank method between groups 1 and 2, in order to confirm the results. The MSTs of patients in groups 1 and 2 were assessed. This analysis did not highlight any statistically significant difference (p>0.05) between the MST of dogs who underwent different cardio-active therapies, and CD as end point.

### Analysis of the groups of treatment in ACVIM class C (2010 to 2018)

Two hundred and two dogs (130 males, 72 females), median age 12.54 years (IQR=3.69), median weight 8.18 Kg (IQR=8.46) were included. One hundred and twenty-one dogs died (59,9%) and 91 (75%) died of CD. The therapy groups and the MST of each dog are reported in Table 4.

**Table 4:** Therapeutic groups and MST of subjects in the ACVIM C class included in the second analysis (2010-2018).

| Therapeutic treatments considered from 2010 to 2018 |  |  |  |  |  |
| --- | --- | --- | --- | --- | --- |
| Groups | No. | % | Therapy | MST | IQR |
| Group 1 | 85 dogs | 42.1 | Furosemide + ACE-I | 318 days | 698 days |
| Group 2 | 76 dogs | 37.6 | Furosemide + ACE-I +<br>Pimobendan | 339 days | 415 days |
| Group 3 | 41 dogs | 20.3 | Furosemide + ACE-I +<br>Pimobendan + Spironolactone | 408 days | 480 days |

The MST was: group 1=318 d (IQR 698), group 2=339 d (IQR 415), and group 3=408 d (IQR 480).

In 70 subjects PH was found, of which 38 (54.3%) were mild, and 32 were moderate-severe (45.7%).

The Kendall tau test revealed a positive correlation between CD and the presence of PH (R=0.2, p=0.005); CD and PH severity (R=0.2, p=0.003); as well as CD and the presence of a moderate-severe PH grade (R=0.18, p=0.011).

The relationship between PH and CD was then investigated in each therapeutic group.

A weak and positive correlation was found in group 1 between the severity of PH and CD (Kendall association coefficient R=0.24, p=0.023), while in groups 2 and 3, the correlation was not statistically significant (R=0.13 and R=0.24, respectively).

The distribution of PH severity among the three groups of dogs who died of CD is reported in Table 5. The Kolmogorov-Smirnov test for independent samples highlighted a statistically significant difference of PH severity distribution only between therapeutic groups 1 and 3 (p=0.033).

**Table 5:** distribution of PH severity in Cardiac Death (CD).

| Distribution of PH severity in CD related to therapeutic class (2001-2018) |  |  |  |  |  |  |
| --- | --- | --- | --- | --- | --- | --- |
| PH severity | Therapeutic Groups |  |  |  |  |  |
|  | 1 |  | 2 |  | 3 |  |
|  | Frequency | % | Frequency | % | Frequency | % |
| Absent | 26 | 63.4 | 16 | 57.1 | 8 | 36.4 |
| Mild | 10 | 24.4 | 7 | 25 | 3 | 13.6 |
| Moderate | 3 | 7.3 | 3 | 10.7 | 6 | 27.3 |
| Severe | 2 | 4.9 | 2 | 7.1 | 5 | 22.7 |
| Total | 41 | 100 | 28 | 100 | 22 | 100 |

The variables considered for survival regression were: sex, therapy, PH severity, weight, age, LA/Ao ratio, E wave, E/A ratio, FE%, FS%, Allo(d) and Allo(s), EDVI, ESVI, and tricuspid regurgitation velocity. Based on the correlation with CD, the chosen predictors were: PH severity, age, LA/Ao ratio, E wave, E/A ratio, Allo(d), and Allo(s). The final model only contained age (HR=1.009, p=0.003) and PH severity (HR=1.316, p=0.032). After adjusting for age, PH severity was a risk factor for CD.

The regression model applied to each group of therapy evidenced different significant correlations. In group 1 the following correlated significantly to CD: E wave, E/A ratio, FS%, Allo(d) and Allo(s) and PH severity. In group 2 the following correlated significantly to CD: LA/Ao ratio, E wave, E/A ratio, EDVI, ESVI and Allo(d) and Allo(s), In group 3 these variables were LA/Ao ratio, EDVI, Allo(d). In groups 1 and 2 considered together as a single group, the variables correlating significantly to CD were LA/Ao ratio, E wave, E/A wave, EDVI, ESVI, Allo(d) and Allo(s) and PH severity.

We found a significant model only in therapeutic group 3, containing predictor LA/Ao (HR=5.867, p=0.014) adjusted for age.

An analysis of the three different endpoints (EP) was performed for each group: CD, first and following therapy changes and moving to more advanced gravity class.

The final step of the survival analysis entailed comparing the clinical findings and the echocardiographic variables at the first and at the last visit for each subject. There was no statistically significant difference between the first and last visit (p>0.05), except for the echocardiographic variable related to the TRV (sign test, p=0.008).

The difference in the tricuspid regurgitation velocity between the first and last visits was the only parameter to be statistically significant (p=0.008) in animals subjected to a therapeutic change in ACVIM C.

## Discussion

DMVD is a progressive disease with a slow onset of clinical signs, and many affected animals die of unrelated diseases [25]. Several studies have reported MSTs and prognostic indicators in dogs with this pathology. However, these studies focused on specific breeds and did not include large breed dogs [26,27,28] or focused on specific aspects of the disease, such as the influence on survival after chordal rupture [29] and the effect of therapy on survival time [30,31]. To the best of our knowledge, there are no studies on the evaluation of MST within the same severity class in relation to the various therapeutic combinations.

The demographic data of the studied population were in line with the data reported in the literature, concerning breed, weight and age [5,25,32].

The literature reports a median survival time of between 5 and 14 months once CHF develops [4,5,30,33]. In our study, the MST between the diagnosis of mitral disease in ACVIM stage C and CD was 531 days (17.7 months) for subjects included from 2001 to 2018, and 335.5 days (11.2 months) for subjects included only from 2010 to 2018. The difference in MSTs between the two populations can be explained by the different classification applied during the study. In veterinary medicine, in order to improve the diagnostic and therapeutic approach to CHF, two classification schemes have been proposed: the ISACH classification and the ACVIM classification [9,16,34]. In this study, it was assumed that for the records included from 2010 to 2018, there was a more standardised classification, not affected by conversion errors.

While the majority of dogs died or were euthanized because of worsening heart failure, multiple factors other than the underlying cardiac disease can impact survival time in veterinary medicine, including medication adherence, financial issues, and owner compliance. Knowledge of MST and prognostic factors could assist clinicians in communicating the prognosis to owners of dogs with advanced heart failure because of DMVD. We believe it is important to understand the long-term outcome and the influence of certain clinical and echocardiographic variables and of the therapeutic scheme on survival in a large series of dogs.

The aim of this retrospective study was to investigate the MST of dogs affected by DMVD belonging ACVIM class C and treated with different combinations of drugs. In addition, the effects of the different therapeutic protocols on the MST and the prognostic values of the echocardiographic data were evaluated.

The clinical records were analysed, the MST was calculated, and the various pharmacological treatments and the changes in class during the follow-up period as well as the prognostic factors were evaluated.

In this study the MST of ACVIM C patients belonging to different therapeutic groups was in accordance with those reported in the literature, although the more complex therapeutic scheme (groups 2 and 3) was associated with a shorter survival time [5,13,35,36,37]. This is despite the fact that patients with more advanced DMVD need more complex cardioactive therapy.

Our study highlighted that with the same severity level of DMVD (subjects in ACVIM class C included from 2001 to 2018), the MST of dogs who died of CD was longer than the MST of those who died of OC. This could mean that cardioactive therapies play a pivotal role in maintaining a good quality of life and in increasing the probability of a longer survival if no other superimposed pathology occurs. Today, DMVD alone is a less frequent cause of death in dogs than in the past.

In line with the literature, we found that LA/Ao and E wave velocity are predictors of CD [35]. The increase in LA/Ao ratio and E-wave values corresponds to an increased risk of CD. The univariate analysis also revealed the E/A ratio, EDVI, ESVI, Allo(d), Allo(s), age and spironolactone administration as predictors of CD. However, only the LA/Ao ratio proved to be significantly correlated in the multivariate analysis, as reported in the literature [35,37].

The univariate analysis of the LA/Ao ratio, E wave, E/A ratio, EDVI, ESVI, Allo(d) and Allo(s) of patients categorized into different therapeutic groups (1, 2 and 3) showed a negative and statistically significant correlation with MST and a significant association with an increased risk of CD. The differences in MSTs among the therapeutic groups were evaluated and an increase in LA/Ao ratio, E wave and EDVI was negatively related to survival.

To the best of our knowledge, no other retrospective study has analysed the MST in different groups of therapy patients belonging to the ACVIM C class. The analysis highlighted that in dogs who died of CD, there was no significant difference in the MST between cardio-active therapy groups, which means that the MST of patients in the ACVIM C class is not related to the therapy group.

The correlation was evaluated between CD and PH in subjects belonging to the ACVIM C class, included from 2010 to 2018, in each therapeutic group. Only in group A was there a positive correlation between severity of PH and risk of CD. The multivariate regression analysis was applied in order to highlight the predictor factors among the clinical and echocardiographic variables, and the uncorrelated variables were selected. Only PH severity and age were positively related to CD. Adjusting for age, the PH severity was shown to be a risk of factor of CD.

The same approach was carried out in each therapy group. Multivariate regression within therapeutic groups showed that only LA/Ao adjusted for age in therapeutic group C was a predictor of CD, and no other references were found in the literature regarding this.

Regarding subjects included from 2010 to 2018, different EPs were considered for each therapeutic group: CD, first and following therapy changes and moving to more advanced severity class. Between the first and last visits, none of the normally distributed variables considered (weight, E wave, E/A ratio, EDVI and Allo(d)) were statistically different. Even for not normally distributed variables (LA/Ao ratio, FS%, ESVI, Allo(s) and TRV), there was no significant difference between the first and last visits, except for TRV. The differences in TRV correlated positively to CD in ACVIM C dogs undergoing therapeutic changes.

As far as the limitations of our study are concerned, this study was performed retrospectively on a population of dogs affected by spontaneous DMVD, and recruited over a long period of time (2001 – 2018), when many changes in diagnostic procedures, therapies and patient classification systems have occurred.

The inclusion criteria of the patients were very strict. However, this is a retrospective study, thus biases cannot be as well controlled as in a well-designed prospective study. Patients who had already been treated with cardioactive therapy were recruited which justifies the variability in therapeutic groups of the overall population.

The echocardiographic values, associated with ACVIM class of DMVD, were useful from a prognostic point of view, and to answer any of the owners’ questions.

The PH severity also correlated strongly to CD and therapeutic groups. In addition, we believe that our study indicates that data regarding therapeutic choices and any variations after the initial diagnosis should be monitored in clinical practice in order to assess the prognosis and modulate the treatment of animals with DMVD.

The retrospective evaluation of the medical records of patients visited over a very long period of time suggests that the classification of DMVD needs revisiting. The ACVIM classification does not include any possible reclassifications into less severe classes of mitral disease, due to the effects of cardioactive therapy.

The lengthening of MST and a good quality of life (QoL) are significant aspects of the therapeutic strategy, and both are very important for the owners. The achievement of a longer MST in our study compared to the literature might be explained by the good compliance of the owners over time, even given the complex protocols [38]. Prospective studies are needed to investigate the compliance effects of owners and the influence of more standardized therapeutic protocols on the QoL and the survival of dogs with DMVD.

